# Epithelial *Ikkβ* deletion modulates immune responses and the IFN_*γ*_/CXCL9 axis during early esophageal carcinogenesis

**DOI:** 10.1101/2025.03.18.643566

**Authors:** Nathan Hodge, Marie-Pier Tétreault

## Abstract

Esophageal cancer is a major cause of cancer-related death, often preceded with chronic inflammation and injuries. The NFκB/IKKβ pathway plays a central role in inflammation, yet its role in early esophageal carcinogenesis remains unclear. This study investigated the role of epithelial IKKβ in early esophageal carcinogenesis. Mice were treated with the carcinogen 4-nitroquinoline-1-oxide (4-NQO) or a vehicle for one month to induce precancerous lesions. Esophagi were harvested and examined through histological, protein, flow cytometry, and RNA analyses. Histological analysis revealed that 4-NQO treatment led to increased inflammation, intraepithelial CD45+ immune cells, and elevated IKKβ phosphorylation levels. Mice with esophageal epithelial-specific *Ikkβ* deletion (4-NQO/*Ikkβ*^*EEC-KO*^) showed delayed progression to a precancerous state, with reduced immune cell recruitment compared to 4-NQO/controls. Immunophenotyping showed decreased recruitment of T cells, including CD4+, CD8+ and regulatory (Tregs) T cells, and increased recruitment of macrophages in 4-NQO/*Ikkβ*^*EEC-KO*^ mice compared to 4-NQO/controls. RNA sequencing data identified 262 differentially expressed genes in 4-NQO/*Ikkβ*^*EEC-KO*^ mice, implicating pathways related to inflammation and wound healing. Notably, the chemokine CXCL9, a T cell chemoattractant, was significantly upregulated in 4-NQO control mice, but not in 4-NQO/*Ikkβ*^*EEC-KO*^ mice. Further analysis identified IFNγ as an upstream regulator of *Cxcl9* expression, and neutralization of IFNγ reduced *Cxcl9* expression levels in 4-NQO treated mice. Additionally, *in vitro* studies demonstrated that IFNγ upregulates *Cxcl9* in an NF-ĸB dependent manner in esophageal keratinocytes. These findings suggest that epithelial IKKβ regulates the immune microenvironment in early esophageal carcinogenesis through the IFNγ/CXCL9 axis and influencing T cell recruitment and inflammatory responses.

**Summary:** In a mouse model of early esophageal squamous cell carcinogenesis, loss of epithelial *Ikkβ* reduced inflammation and T cell recruitment, increased macrophage recruitment, inhibited IFNγ/CXCL9 signaling, and delayed the transition to a precancerous state.

## Introduction

Esophageal cancer (EC) has a 5-year survival rate of less than 20%, making it the 6^th^ leading cause of cancer death worldwide [1,2]. This high mortality rate is due to most patients being diagnosed at a late stage [1,3]. Squamous cell cancer, the most aggressive form of EC, accounts for over 90% of cases [1,3]. The pathogenesis of ESCC involves a multi-step process from inflammation, hyperplasia, and dysplasia to carcinoma in situ to invasive carcinoma [3,4]. Chronic inflammation, caused by irritating factors such as alcohol, tobacco and hot beverages, plays a significant role [1,3]. The late diagnosis of ESCC has hindered understanding of the molecular mechanisms of the early stages of the disease. Therefore, a better understanding of these mechanisms is urgently needed to improve diagnosis and treatment options for this deadly cancer.

The epithelial tissue is primarily affected by chronic inflammation in the early stages of esophageal carcinogenesis, which significantly contributes to cancer development [5,6]. This inflammatory environment can alter the immune landscape, profoundly impacting tumor development [7]. Epithelial cells play a central role in shaping the tissue response during the early inflammatory phase of carcinogenesis by releasing cytokines and chemokines [8]. These signaling molecules attract specific immune cell subsets to the site of inflammation, thereby guiding the immune response [8]. Thus, the crosstalk between epithelial and immune cells is crucial in this context, as it influences the progression of the disease.

The IKKβ/NF-κB pathway is a key player in the crosstalk between epithelial and immune cells, acting as a central signaling hub in inflammatory responses [9-11]. This pathway is activated upon local tissue damage or inflammatory cytokine signaling [9-11]. IKKβ activates canonical NFκB signaling by phosphorylating the inhibitory molecule IκB, resulting in its polyubiquitination and proteasomal degradation. This process releases NF-κB dimers, which then translocate to the nucleus to induce the transcription of target genes [12]. The list of IKKβ/NFκB target genes is extensive and includes those involved in inflammation, immunity, stress responses, angiogenesis, cell survival, and proliferation [13]. The role of IKKβ signaling varies depending on the tissue, cell type, and disease state [14-16]. A pro-tumorigenic role for IKKβ has been demonstrated in several mouse models, including those of intestinal, pancreatic, and liver cancer[17-20]. In addition to being dysregulated in ESCC, we recently observed that IKKβ/NFκB signaling is elevated in inflamed epithelial tissue adjacent to esophageal tumors [21]. However, the specific role of epithelial IKKβ/NFκB signaling in the transition from a normal state to the early precancerous stage of ESCC remains unknown. In this study, we investigated how IKKβ signaling in esophageal epithelial cells regulates the inflammatory response during the early precancerous stage of esophageal squamous cell carcinogenesis.

## Materials and Methods

### Generation of ED-L2-Cre/Ikkβ^L/L^ Mice

*ED-L2-Cre/Ikkβ*^*L/*L^ mice (ref: *Ikkβ*^*EEC-KO*^) were generated by crossing *Ikkβ* floxed mice [22] with mice harboring *Cre* recombinase under the control of the *EBV-ED-L2* promoter [23]. All mice used for experiments were on a pure c57BL/6 background. Sex-matched littermate *Ikkβ*^LoxP/LoxP^ mice (ref: control) were used as controls. Both male and female experimental groups were included. Mouse esophagi were harvested and processed for histology. Tissue was fixed in neutral buffered formalin (Fisher Scientific, Pittsburgh, PA) for 24 hours, embedded in paraffin, and 4-μm sections were applied to charged plus slides. Esophageal sections were also stained with hematoxylin and eosin for histological examinations. The following numbers of matched littermate control and *Ikkβ*^*EEC-KO*^ mice were examined histologically: 1 month 4-NQO or Propylene glycol (vehicle) treatment: 6 controls, 6 *Ikkβ*^*EEC-KO*^, 6 wild-type c57/BL6. 4 month 4-NQO or vehicle treatment; 5 controls and 5 mutants; 4-NQO treated mice: 5 controls and 5 mutants.

### 4-NQO mouse treatment

Treatment with the carcinogen 4-NQO (Sigma, St. Louis, MO) was performed on 8 week-old mice. 4-NQO (100μg/ml) or 2% propylene glycol (vehicle) (Fisher Scientific) was administered in drinking water for 1 month or 4 months to induce different stages of esophageal squamous cell carcinogenesis. Mice were weighed weekly and body condition was scored and recorded.

### Primary cell cultures and treatment

Primary cultures of mouse esophageal epithelial cells were generated from 8 week-old c57 BL/6 mice treated for 1 month with 2% propylene glycol or 100μg/ml 4-NQO. Primary cultures did not get converted into cell lines. Briefly, the mouse esophagus was harvested and treated with 20% dispase (Corning, Corning, NY) at 37°C for 10 minutes. The epithelium was mechanically separated and treated with 0.25% trypsin at 37°C for 10 minutes to generate single cell suspensions. Primary cultures were then cultured in K-SFM supplemented with 40 μg/ml bovine pituitary extract, 1.0 ng/ml EGF, and penicillin/streptomycin (Thermo Fisher, Pittsburg, PA). Primary cultures of esophageal epithelial cells were treated with 50ng/mL recombinant IFNγ (R&D Systems, Minneapolis, MN), with or without 10μg/mL JSH23 (Sigma) for 4h at 37°C.

### Immunohistochemistry/Immunofluorescence

Heat antigen-retrieval (2100 Antigen Retriever, Electron Microscopy Sciences, Hatfield, PA) was performed on formalin-fixed paraffin-embedded esophageal sections, and slides were incubated with the following primary antibodies: 1:100 goat anti-CXCL9 (#AF-492-NA, R&D Systems), 1:500 rabbit CD45 (#AB10558, Abcam, Cambridge, MA), 1:50 rabbit anti-CD3 (#78588, Cell Signaling, Danvers, MA), 1:200 rabbit anti-CD4 (#183685, Abcam), 1:50 rabbit anti-CD8 (#98941, Cell Signaling), 1:50 rat anti-FOXP3 (#14-5773-82, Invitrogen, Pittsburgh, PA), 1:50 rabbit anti-F4/80 (#70076, Cell Signaling). Species-specific secondary antibodies were added, and detection was performed as previously described[15]. For fluorescent labeling, Alexa-conjugated antibodies (Thermo Fisher Scientific) were used. Images were captured using a Nikon Eclipse Ci-E microscope with a Nikon DS-Ri2 camera and NIS Elements software. Quantification was performed by determining the number of immune cell positive counts per high power field.

### Western Blots and antibody arrays

Mouse esophageal epithelia were homogenized in triton lysis buffer (1% Triton X-100, 50mM Tris-HCl pH 7.5, 100mM NaCl, 5mM EDTA, 40mM β-glycerophosphate, 5% glycerol, 50mM NaF) plus protease (Pierce, Rockford, IL) and phosphatase inhibitors (Sigma-Aldrich). Pierce BCA protein assay was used to determine protein concentration (Thermo Scientific). Protein separation was done on Novex NuPage 4–12% gels (Thermo Scientific) and transferred onto polyvinylidene difluoride membrane (EMD Millipore, Billerica, MA). After blocking in 5% non-fat dry milk, membranes were incubated overnight at 4°C with the following antibodies: 1:750 rabbit anti-phosphorylated IKKα/β Ser^176/180^ (Cell Signaling), 1:3000 rabbit anti-IKKβ (Cell signaling), 1:10,000 Rabbit anti-GAPDH (Sigma). The concentration of 200 different mouse cytokines in lysates of mouse esophageal epithelial cells was determined by using the mouse Quantibody Array 4000 testing service (RayBiotech, Inc., Norcross, GA).

### RNA analyses

RNA was extracted using the RNeasy mini purification kit (Qiagen, Germantown, MD) and Reverse transcription was performed with the Maxima First-Strand cDNA Synthesis for RT-qPCR kit (Thermo Fisher Scientific). Quantitative real-time PCR was performed using Taqman Universal Master Mix (Applied Biosystems, Pittsburg, PA). The TATA box binding protein or Glyceraldehyde-3-phosphate dehydrogenase genes were used as the internal control.

### RNA sequencing

For RNA sequencing, cDNA libraries were generated using the Tru-Seq Total RNA-seq library prep (Illumina, San Diego, CA) following the manufacturer’s instructions. Sequencing was performed using Illumina HiSeq 4000 (Northwestern University NUSeq Core Facility). The quality of DNA reads, in fastq format, was evaluated using FastQC. Adapters were trimmed and reads of poor quality or aligning to rRNA sequences were filtered. The cleaned reads were aligned to the *Mus musculus* genome (mm10) using STAR [24]. Htseq-count [25] was used in conjunction with a gene annotation file for mm10 obtained from UCSC (University of California Santa Cruz; http://genome.ucsc.edu) to calculate read counts for each gene. DESeq2 [26] was used to determine normalization and differential expression. The cutoff for determining significantly differentially expressed genes was an FDR-adjusted p-value less than 0.05. The Metascape online platform (http://metascape.org/) [27] was used to identify gene ontology (GO) terms.

### Flow Cytometry

For flow cytometry experiments, mouse esophagi were harvested. The esophageal epithelium was mechanically separated from whole esophageal tissue, and both tissue fractions were minced. Esophageal epithelia were digested with 0.05% Trypsin-EDTA (Thermo Fisher Scientific) for 2 minutes at 37°C, while the remaining esophageal tissue was digested with collagenase V (Sigma-Aldrich) for 15 minutes at 37°C. Esophageal cell suspensions were filtered using 70 μm nylon filters. Up to 1 million cells was stained for each sample. Live/dead cell stain was performed using eBioscience™ Fixable Viability Dye eFluor™ 506 (Thermo Fisher Scientific) according to the manufacturer’s instructions. Cell surface staining was performed for 30 minutes using the indicated antibodies. Intracellular staining was performed using a Foxp3 fixation/permeabilization kit (BioLegend, San Diego, CA) according to manufacturer’s instructions. T cells were phenotyped as CD45^+^CD11b^−^Ly6G^−^CD19^−^CD3^+^, B cells as CD45^+^CD11b^−^Ly6G^−^CD19^+^CD3^−^, monocytes as CD45^+^CD115^+^, neutrophils as CD45^+^CD11b^+^Ly6G^+^, macrophages as CD45^+^F4/80^+^, eosinophils as CD45^+^SiglecF^+^, dendritic cells as CD45^+^CD11c^+^, mast cells as CD45^+^CD117^+^, natural killer cells as CD45^+^,CD49b^+^,NK1.1^+^, regulatory T cells as CD45^+^CD11b^−^Ly6G^−^CD19^−^CD3^+^CD4^+^CD8^−^Foxp3^+^, cytotoxic T lymphocytes as CD45^+^CD11b^−^Ly6G^−^CD19^−^CD3^+^CD8^+^,CD4^−^ and T-helper cells as CD45^+^CD11b^−^Ly6G^−^CD19^−^CD3^+^CD4^+^,CD8^−^. Flow cytometry experiments were run on a BD FACSymphony A5 (BD Biosciences, San Jose, CA) and data was analyzed using FlowJo (BD Biosciences).

### Antibodies for flow cytometry

The table below lists the antibodies for flow cytometry experiments.

| Antibody | Clone | Fluorophore | Company | Catalog Number |
| --- | --- | --- | --- | --- |
| CD115 | T38-320 | BV421 | BD Biosciences | 743638 |
| CD117 | 2B8 | BV711 | BD Biosciences | 563160 |
| CD11b | M1/70 | BUV737 | BD Biosciences | 564443 |
| CD11c | HL3 | BUV395 | BD Biosciences | 564080 |
| CD170 | E50-2440 | PE-CF594 | BD Biosciences | 562757 |
| CD19 | 1D3 | APC-Cy7 | BD Biosciences | 557655 |
| CD31 | 390 | AF488 | BD Biosciences | 563607 |
| CD3e | 500A2 | AF700 | BD Biosciences | 557984 |
| CD4 | GK1.5 | BUV496 | BD Biosciences | 564667 |
| CD44 | 1M7 | BV786 | BD Biosciences | 563736 |
| CD45.2 | 104 | BV650 | BD Biosciences | 740490 |
| CD49b | HMa2 | BV605 | BD Biosciences | 740363 |
| CD8a | 53-6.7 | BB700 | BD Biosciences | 566409 |
| F4/80 | T45-2342 | A647 | BD Biosciences | 565854 |
| FoxP3 | FJK-16s | PE | eBioscience | 12-5773-80 |
| Live/Dead | N/A | 506e | eBioscience | 65-0866-14 |
| Ly6G | 1A8 | PE-Cy7 | BD Biosciences | 560601 |
| NK 1.1 | PK136 | BB630 | BD Biosciences | 624294 |

### In vivo IFNγ-neutralizing treatment

For the last 1.5 weeks of 4-NQO treatment, mice were injected twice a week with 200 μg of either anti-mouse IFNγ (#BE0055, BioXCell, West Hanover, NH) or rat IgG1 Isotype control (#BE0290, BioXCell) for a total of three injections. Mice were harvested the day after the last injection.

### Statistical analyses

Results were expressed as mean ± standard error of the mean, with statistical significance of differences between experimental conditions established at 95%. One-way analysis of variance (ANOVA) followed by the post-hoc Tukey test were used to compare the statistical difference between means of all groups. The Sidak multiple comparison test was used to compare the statistical difference between means of preselected groups. Pairwise comparisons were compared using Student’s t test to determine the statistical difference. All stats were done using GraphPad Prism version 4.0 (GraphPad Software, San Diego, CA).

## Results

### IKKβ signaling is activated in precancerous esophageal epithelium in vivo

Increased activation of IKKβ/NFκB signaling has been reported in patients with esophageal chronic inflammation and hyperplasia of the esophagus, two precursors to ESCC development [21]. However, the limited availability of early-stage patient samples restricts the study of IKKβ/NFκB signaling in esophageal carcinogenesis. A well-established mouse model of ESCC involves administering the carcinogen 4-NQO in drinking water for four months, followed by three months of regular water [28]. Recently, this model has been adapted to study molecular changes in early esophageal carcinogenesis using shorter 4-NQO treatments [29,30]. To investigate the role of IKKβ/NFκB signaling in these early stages, we first evaluated the phosphorylation levels of IKKβ in mice treated with 4-NQO or vehicle control for one month (**Figure 1A**). Histological analysis revealed inflamed epithelial tissue with large pockets of immune cells and basal cell hyperplasia (**Figure 1B**). Increased IKKβ phosphorylation levels were observed in esophageal epithelial enrichments from one month treated 4-NQO mice compared with vehicle treated mice (**Figure 1C-D**).

**Figure 1.**
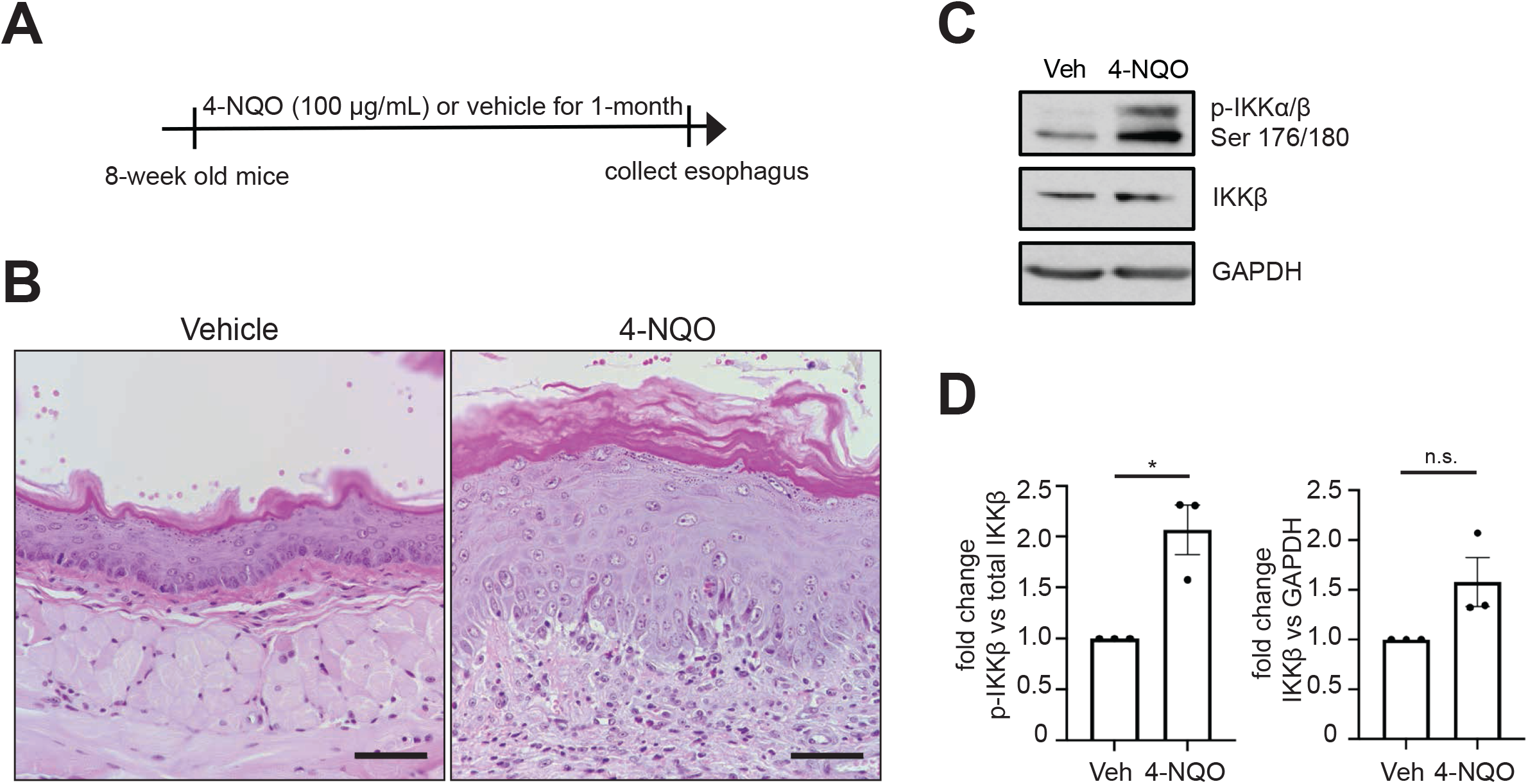
Increased IKKβ signaling is detected in the esophageal mucosa of mice treated with 4-NQO. (**A**) Schematic of 4-NQO treatment. (**B**) H&E staining of esophageal sections of mice treated with vehicle or 4-NQO for one month. n = 3, scale bar = 50 µM. (**C**) Western blot showing P-IKKα/β^Ser176/180^ and IKKβ expression in the esophageal mucosa of mice treated with vehicle or 4-NQO for one month. GAPDH was used as a loading control. n = 3. (**D**) Densitometric analysis of the ratio of phosphorylated IKKβ to total IKKβ protein and of total IKKβ to GAPDH protein. n=3. Two-tailed paired T test, *P ≤ 0.05. Bar graph represents mean ± SEM.

### Loss of esophageal epithelial Ikkβ delays the transition from a normal to a precancerous inflammatory state

To determine the role of epithelial IKKβ signaling in early esophageal carcinogenesis, we generated mice with esophageal epithelial *Ikkβ* loss (*Ikkβ*^*EEC-KO*^ mice) by crossing *Ikkβ* floxed mice [22] with *ED-L2/Cre* mice [23]. *Ikkβ*^*EEC-KO*^ mice and their littermate controls were treated with 4-NQO or vehicle for one month (**Figure 2A**). Decreased IKKβ expression in *Ikkβ*^*EEC-KO*^ mice were confirmed by Western blot (**Figure 2B-C**). Vehicle/*Ikkβ*^*EEC-KO*^ mice showed no histological changes compared to littermate controls (**Figure 2D**). 4-NQO/control mice exhibited esophageal basal cell hyperplasia and pockets of recruited immune cells, whereas 4-NQO/*Ikkβ*^*EEC-KO*^ mice showed a notable decrease in immune cell recruitment (**Figure 2D**). To further investigate the impact of *Ikkβ* loss during the transition from a normal to a precancerous state, *Ikkβ*^*EEC-KO*^ mice and their littermate controls were treated with 4-NQO or vehicle for four months (**Figure 2E**). Vehicle/*Ikkβ*^*EEC-KO*^ mice showed no histological changes compared to vehicle/control mice (**Figure 2E**). However, 4-NQO treatment led to significant disruption of the epithelial layer and an increase in immune cell recruitment in control mice, which was greatly attenuated in *Ikkβ*^*EEC-KO*^ mice (**Figure 2E**). CD45 immunostaining further confirmed the decreased recruitment of immune cells in 4-NQO/*Ikkβ*^*EEC-KO*^ mice compared with 4-NQO/controls (**Figures 2F-G**).

**Figure 2.**
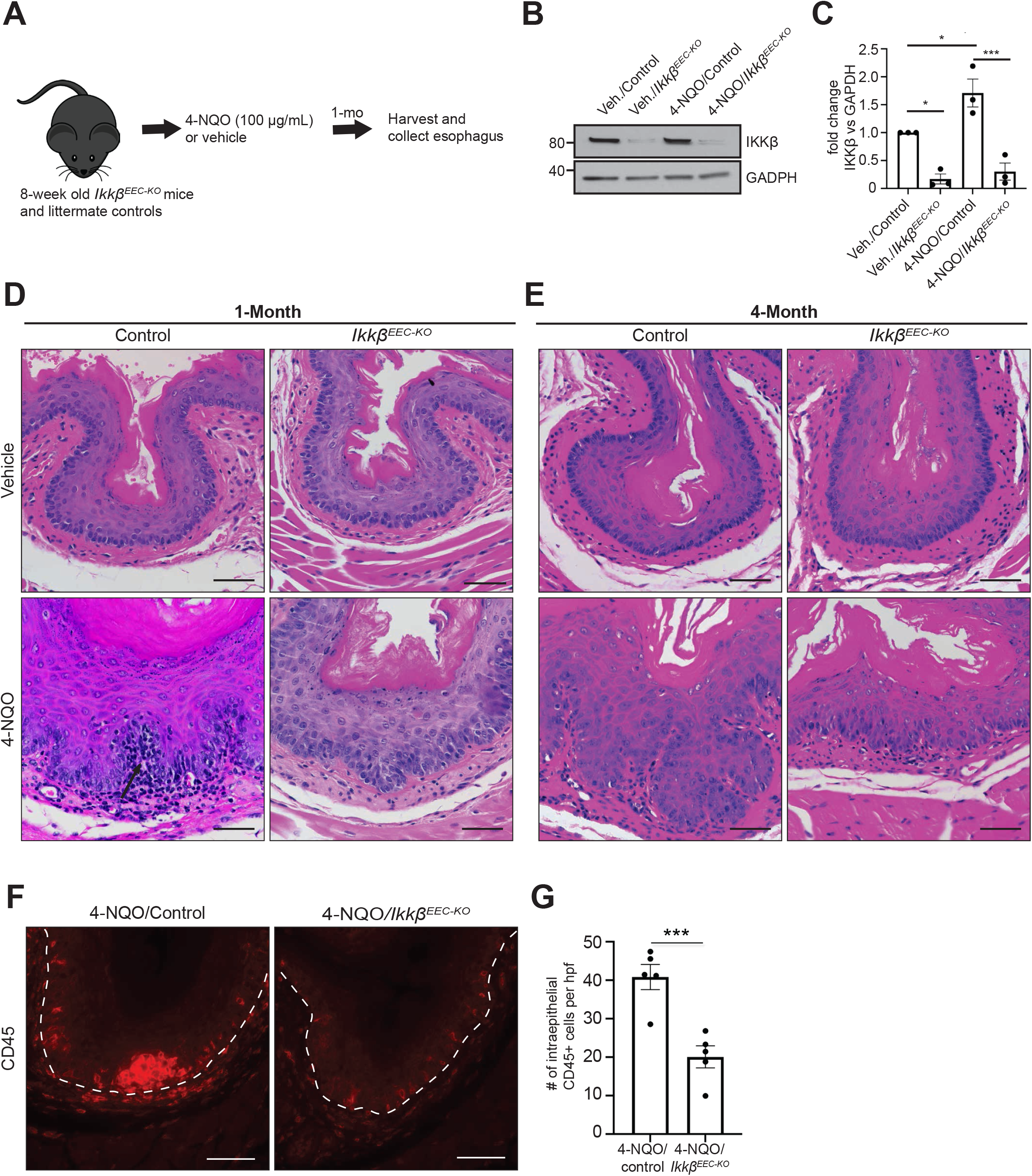
Epithelial *Ikkβ* deletion in the esophagus delays the transition from a normal to a precancerous inflammatory state. (**A**) Schematic of 4-NQO treatment in *Ikkβ*^*EEC-KO*^ mice and littermate controls. (**B**) Western blot of IKKβ expression in the esophageal mucosa of *Ikkβ*^*EEC-KO*^ mice and littermate controls treated with vehicle (PG) or 4-NQO for one month. GAPDH was used as a loading control. n = 3 (**C**) Densitometric analysis of the ratio of total IKKβ to GAPDH protein. n = 3. One way ANOVA followed by a Post-Hoc Tukey Test, *P ≤ 0.05, ***P ≤ 0.001. Bar graph represents mean ± SEM. (**D-E**) H&E of esophageal sections of *Ikkβ*^*EEC-KO*^ mice and littermate controls treated with vehicle (PG) or 4-NQO for one month (**D**) or 4 months (**E**). n = 6, scale bar = 50 µM. Arrow is indicating a pocket of immune cells. (**F**) CD45 immunofluorescence staining of esophageal sections of *Ikkβ*^*EEC-KO*^ mice and littermate controls treated 4-NQO for one month. n = 5, scale bar = 50 µM. (**G**) Bar plot showing the mean of CD45+ counts per hpf ± SEM. n = 5. Paired T test, ***P ≤ 0.001.

### Transcriptomics analysis identifies inflammatory pathways differentially regulated by Ikkβ during the transition from normal to precancerous condition

To identify potential targets of *Ikkβ* in 4-NQO-induced esophageal inflammation and pre-cancer, we analyzed gene expression changes by performing RNA sequencing on *Ikkβ*^*EEC-KO*^ mice and littermate controls treated with 4-NQO for one month. Overall, 262 genes were significantly differentially regulated (false-discovery rate-adjusted *P* value <0.05, |log2 fold change| ≥ 1.0). Of these, 92 genes were upregulated and 170 were downregulated (**Figure 3A, Supplementary Table 1**). The gene expression profile was visualized using a volcano plot (**Figure 3B**). To determine which pathways were differentially regulated following the loss of epithelial *Ikkβ*, we performed gene set enrichment of GO terms. As shown in **Figure 3C**, changes in gene regulation were observed in pathways related to response to keratinocyte differentiation, response to wounding, cell chemotaxis, positive regulation of T cell differentiation in thymus, defense response to bacterium, negative regulation of T cell apoptosis and antimicrobial humoral response (**Figure 3C**).

**Figure 3.**
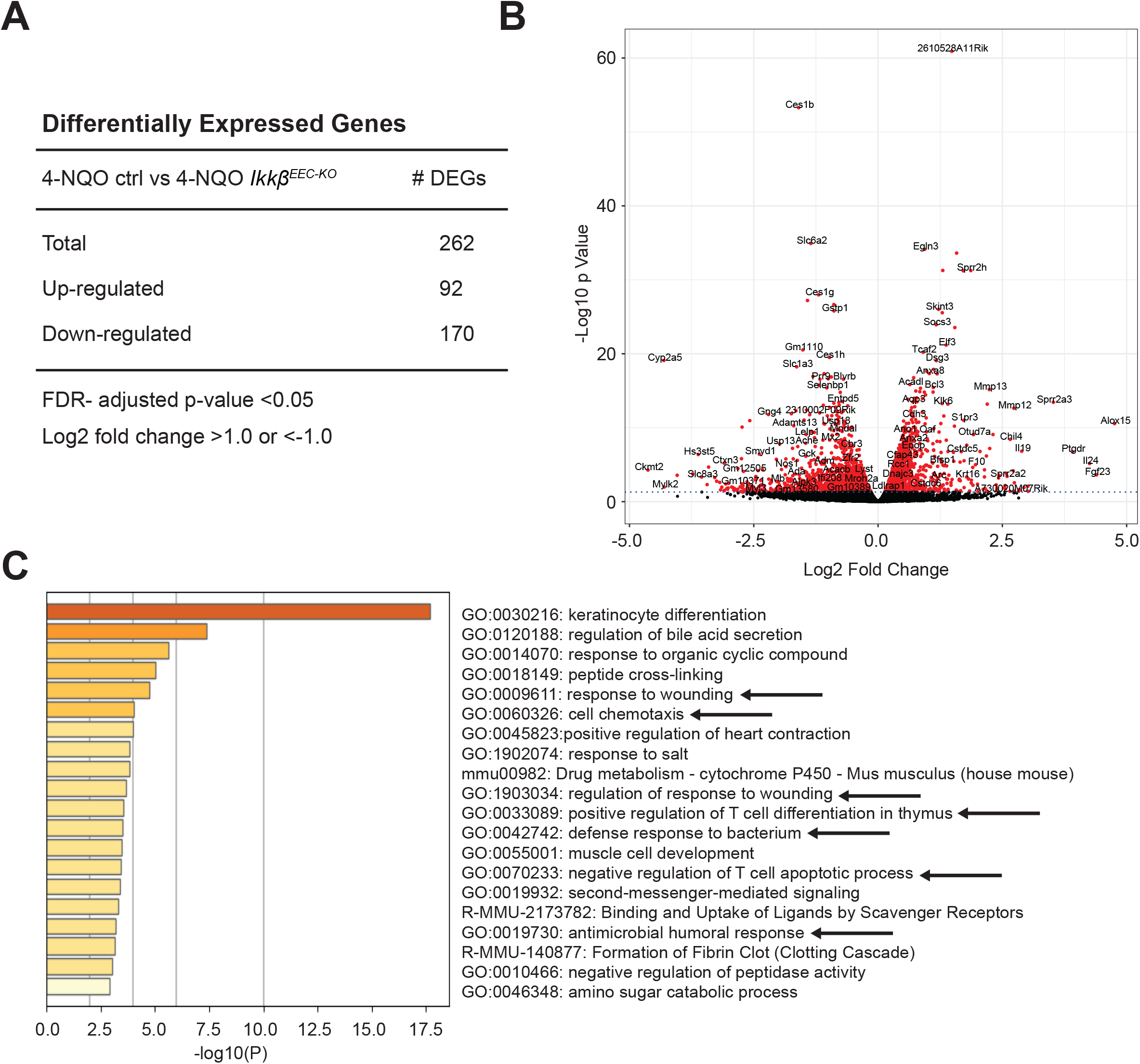
Transcriptomics analysis identifies inflammatory pathways differentially regulated by *Ikkβ* during the transition from normal to precancerous condition. (**A**) Number of differentially expressed genes (DEGs) in 4-NQO/*Ikkβ*^*EEC-KO*^ mice compared to 4-NQO/controls based on a log2 fold change of 1.0 and FDR ≤ 0.05. n = 5. (**B**) Volcano plot showing the log2 fold change in gene expression and the statistical significance. (**C**) Gene Ontology (GO) identified enriched functional pathways using DEGs from 4-NQO/*Ikkβ*^*EEC-KO*^ mice and 4-NQO/controls.

### Loss of epithelial Ikkβ leads to an increase in macrophage recruitment in early esophageal carcinogenesis

CD45 immunostaining showed significant changes in immune cell recruitment and several of the top enriched pathways indicated changes in inflammatory and wound healing-related genes. To characterize the esophageal immune landscape in 4-NQO/control and 4-NQO/*Ikkβ*^*EEC-KO*^ mice, we performed flow. Interestingly, F4/80+ macrophages were significantly enriched in the esophagus of 4-NQO/*Ikkβ*^*EEC-KO*^ mice compared to 4-NQO/controls (**Figure 4A**). Immunostaining for F4/80 confirmed an increased presence of intraepithelial F4/80+ macrophages in 4-NQO/*Ikkβ*^*EEC-KO*^ mice (**Figure 4B-C**). Macrophages are known to play a role in wound repair [31]. As shown in **Figure 4D**, genes involved in wound healing, including *Alox15, Mmp12, F10, F7, HB-EGF, Krt6a, Anxa8*, many of which are associated with or produced by macrophages [32-35] had increased expression in 4-NQO/*Ikkβ*^*EEC-KO*^ mice by RNA-sequencing. Quantitative PCR confirmed increased mRNA expression levels of *Alox15* and *Mmp12* in 4-NQO/*Ikkβ*^*EEC-KO*^ mice compared to 4-NQO/control mice (**Figure 4E**).

**Figure 4.**
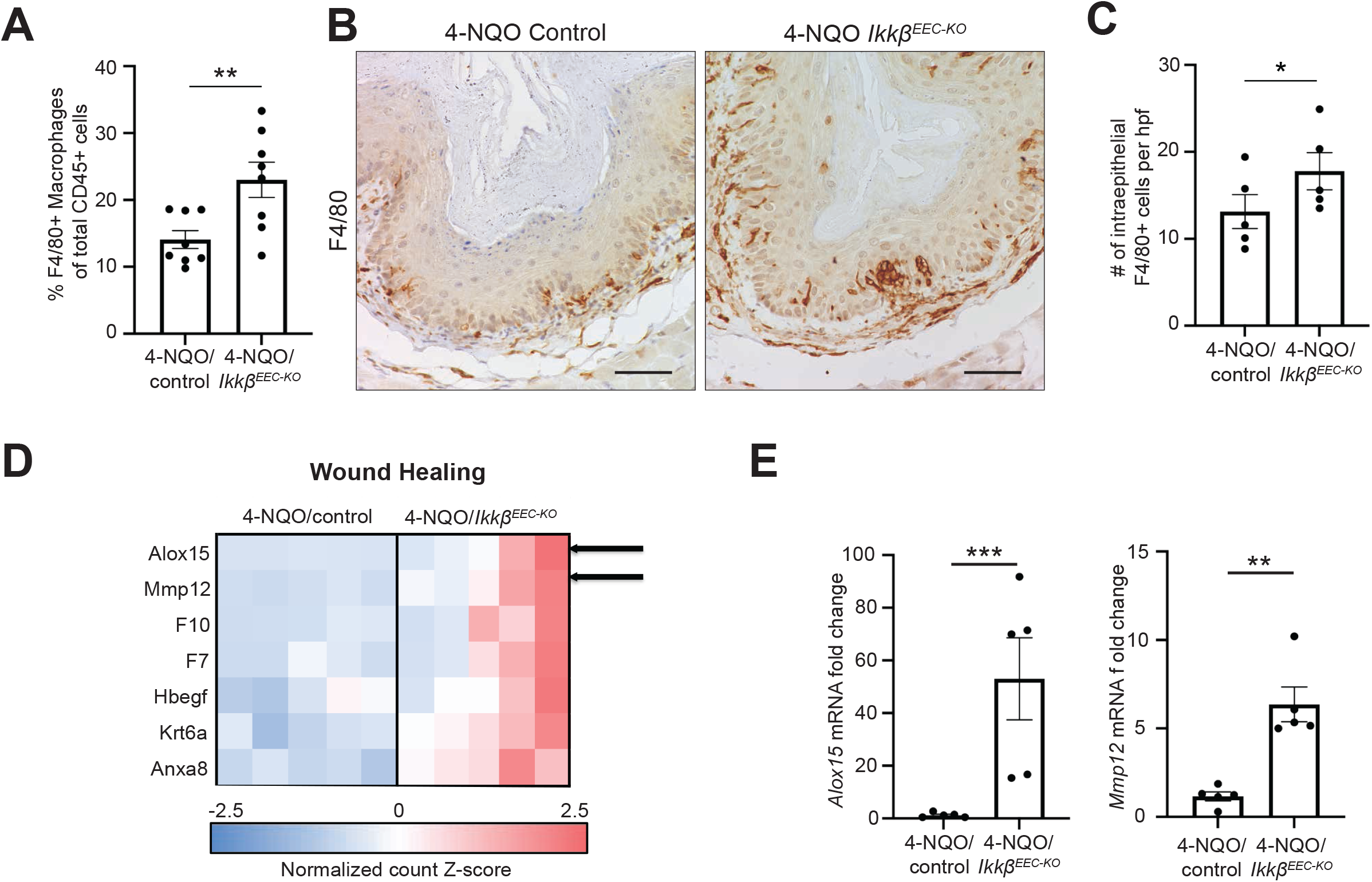
Loss of epithelial *Ikkβ* leads to an increase in macrophage recruitment to the esophagus during early esophageal carcinogenesis. (**A**) Flow cytometry analysis showing the percentage of F4/80+ macrophages per total CD45+ cells. n = 8. Two-tailed paired T test, **P ≤ 0.01. (**B**) F4/80 immunohistochemistry of esophageal sections of 4-NQO/control and 4-NQO/*Ikkβ*^*EEC-KO*^ mice. n = 5. Scale bar = 50 µM. (**C**) Bar plot showing the mean of F4/80+ counts per hpf ± SEM. n = 5. Two-tailed paired T test, *P ≤ 0.05. (**D**) Heatmap showing normalized expression values of differently expressed genes involved in wound healing. n = 5. (**E**) *Alox15* and *Mmp12* mRNA expression levels were determined in the esophageal mucosa of 4-NQO/control and 4-NQO/*Ikkβ*^*EEC-KO*^ mice using qPCR. n = 5. Bar graphs represent mean ± SEM. Two-tailed paired T test, **P ≤ 0.01, ***P ≤ 0.001.

### T cell recruitment is dependent on epithelial IKKβ in the transition to a precancerous state

In addition to the increased recruitment of intraepithelial macrophages, we also detected a significant decrease in the recruitment of CD3+, CD4+, and CD8+ T cells by flow cytometry in 4-NQO/*Ikkβ*^*EEC-KO*^ mice compared to 4-NQO/control mice (**Figure 5A-C**), while no change was observed in the number of Tregs (**Figure 5D**). Immunostaining confirmed a significant decrease in the recruitment of intraepithelial CD3+ T cells and CD4+ T cells to the esophagus of 4-NQO/*Ikkβ*^*EEC-KO*^ mice compared to 4-NQO/control mice (**Figure 5E-H)**. We were unable to confirm a significant increase in the recruitment of CD8+ T cells by immunostaining due to the low number of cells detected (**Figure 5I-J**). Furthermore, although we observed no significant change in the number of Tregs by flow cytometry (**Figure 5D**), we detected a significant decrease in intraepithelial Tregs by immunostaining (**Figure 5K-L**). Further supporting the observation of decreased recruitment of T cells, our gene enrichment analysis showed a decrease in the expression of genes related to leukocyte migration (**Figure 5M**). Among the genes differentially expressed and linked to T cell migration were the chemoattractant *Cxcl9* and its receptor Cxcr3 [36]. Cxcr3 is known to be expressed on CD8+ T cells and Th1 CD4+ T cells [36]. This suggests that IKKβ regulates the CXCL9/CXCR3 axis in T cell recruitment during the transition from normal to precancerous state in the esophagus.

**Figure 5.**
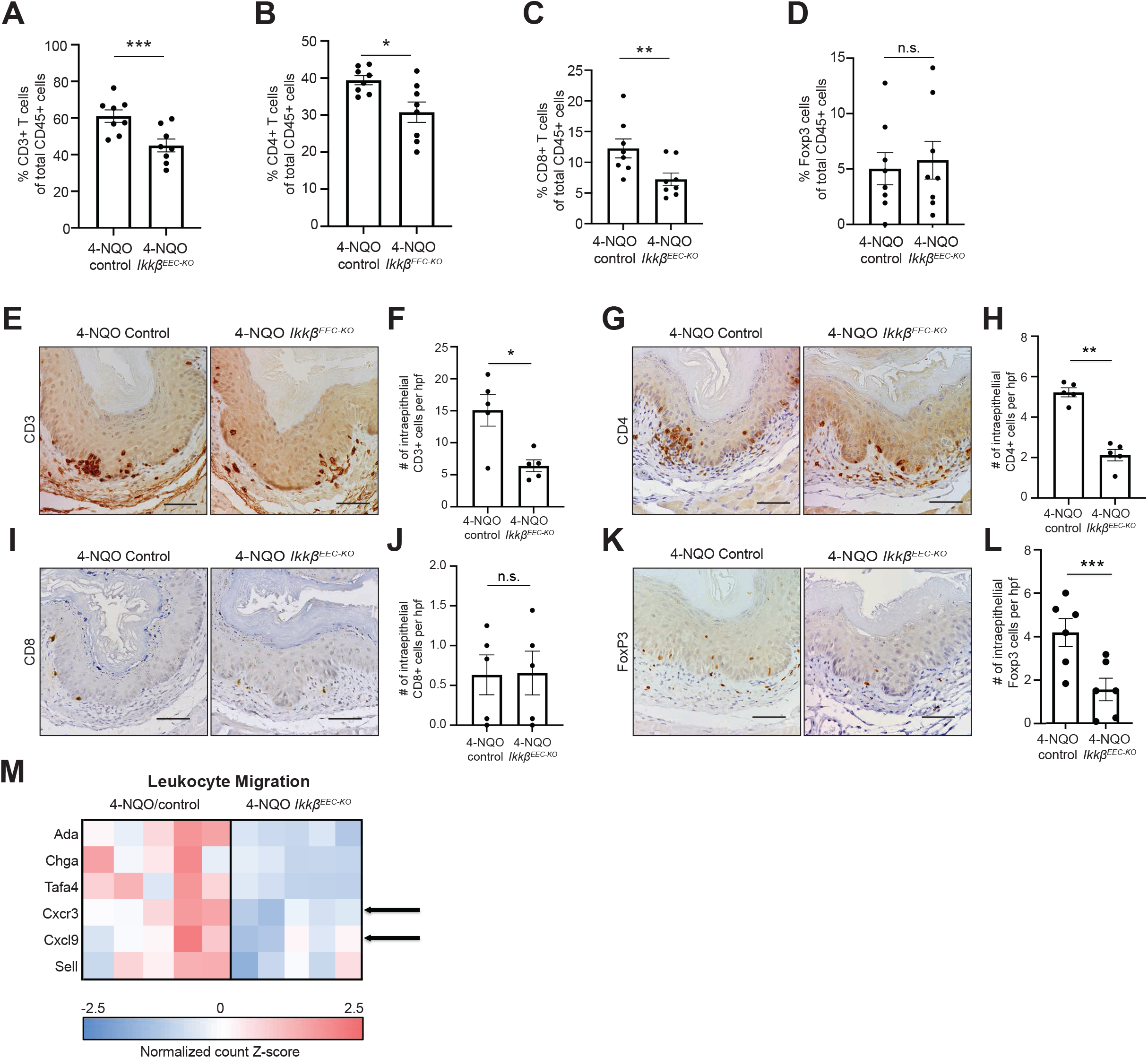
Epithelial IKKβ signaling regulates T cell recruitment during the transition from normal to a precancerous state. (**A-D**) Flow cytometry analysis showing the percentage of CD3+ T cells (**A**), CD4+ T cells (**B**), CD8+ T cells(**C**), or Tregs (**D**) per total CD45+ cells. n = 8. Two-tailed paired T test, *P ≤ 0.05, **P ≤ 0.01. (**E**) CD3 immunohistochemistry of esophageal sections of 4-NQO/control and 4-NQO/*Ikkβ*^*EEC-KO*^ mice. n = 5. Scale bar = 50 µM. (**F**) Bar plot showing the mean of CD3+ counts per hpf ± SEM. n = 5. Two-tailed paired T test, *P ≤ 0.05. (**G**) CD4 immunohistochemistry of esophageal sections of 4-NQO/control and 4-NQO/*Ikkβ*^*EEC-KO*^ mice. n = 5. Scale bar = 50 µM. (**H**) Bar plot showing the mean of CD4+ counts per hpf ± SEM. n = 5. Two-tailed paired T test, **P ≤ 0.01, (**I**) CD8 immunohistochemistry of esophageal sections of 4-NQO/control and 4-NQO/*Ikkβ*^*EEC-KO*^ mice. n = 5. Scale bar = 50 µM. (**J**) Bar plot showing the mean of CD8+ counts per hpf ± SEM. (**K**) Foxp3 immunohistochemistry of esophageal sections of 4-NQO/control and 4-NQO/*Ikkβ*^*EEC-KO*^ mice. n = 6. Scale bar = 50 µM. (**L**) Bar plot showing the mean of Foxp3+ counts per hpf ± SEM. Two-tailed paired T test, ***P ≤ 0.001. (**M**) Heatmap showing normalized expression values of differently expressed genes involved in leukocyte migration. n = 5.

### Epithelial IKKβ leads to production of CXCL9 through IFNγ signaling

CXCL9 is a known T-cell chemoattractant and has been shown to be expressed in epithelial cells [36-38]. We next sought to investigate the role of epithelial IKKβ signaling in the downstream regulation of CXCL9 signaling. We first analyzed CXCL9 expression levels in esophageal epithelia of mice by multiplex ELISA (**Figure 6A**). Compared to vehicle treated mice, we observed a significant increase in CXCL9 expression in 4-NQO/control mice, and this increase was significantly attenuated in 4-NQO/*Ikkβ*^*EEC-KO*^ mice (**Figure 6A**). To confirm that epithelial cells were the source of CXCL9 and to visualize their potential association with recruited CD3+ T cells, we performed co-staining for CXCL9 and CD3. We observed that CXCL9 was detected in distinct cell populations than the CD3+ T cells in the epithelial tissue (**Figure 6B**).

**Figure 6.**
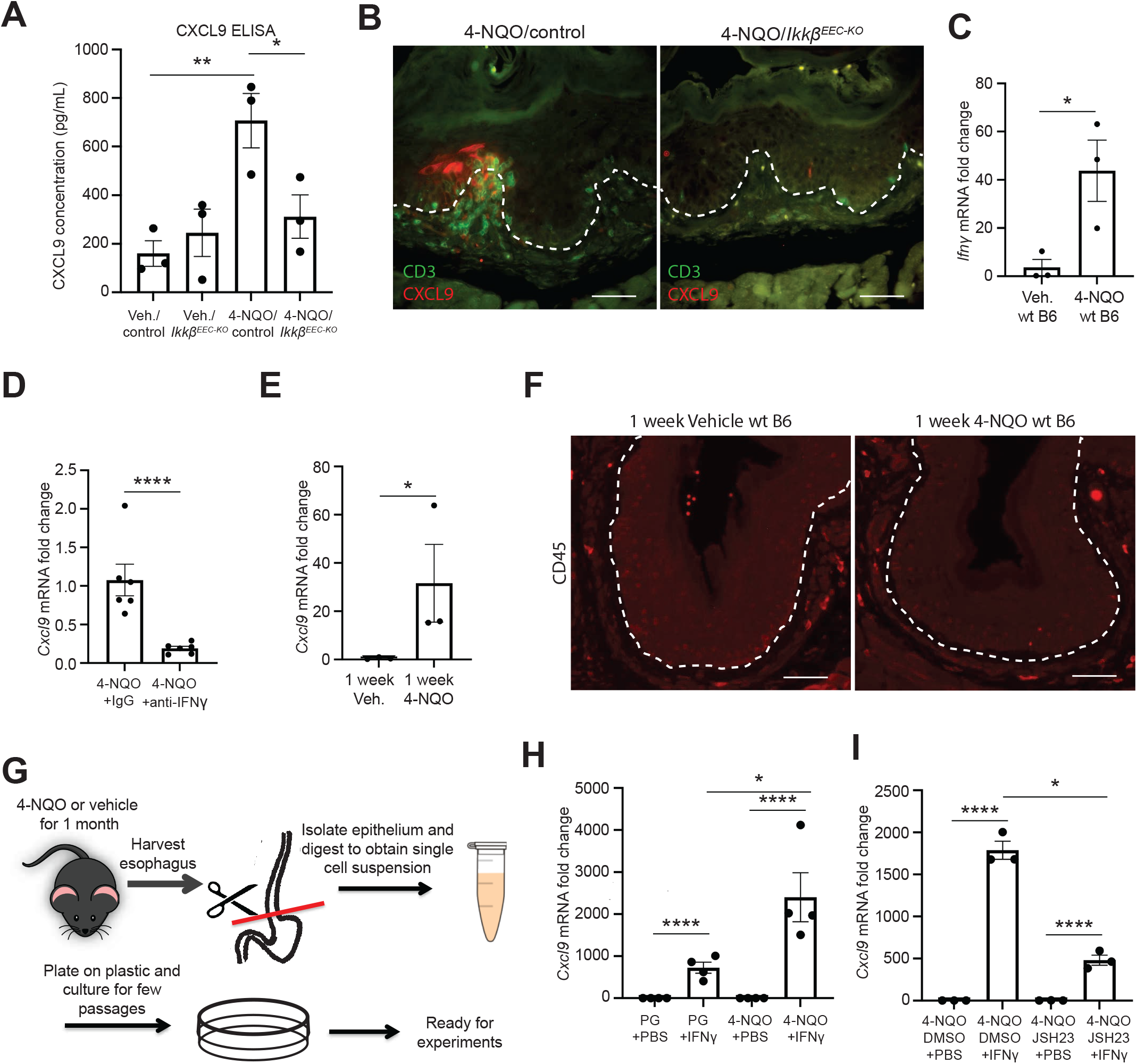
*Cxcl9* expression is regulated through epithelial IKKβ/NFκB signaling and IFNy. (**A**) Multiplex ELISA show changes in CXCL9 production in control and *Ikkβ*^*EEC-KO*^ mice treated with vehicle (PG) or 4-NQO for one month. n = 3. One-way analysis of variance followed by a Sidak multiple comparisons test, *P ≤ 0.05, **P ≤ 0.01. (**B**) CXCL9 (red) and CD3 (green) co-immunofluorescence of esophageal sections of 4-NQO mice. n = 6. Scale bar = 50 µM. (**C**) Evaluation of *Ifnγ* expression levels in mice treated with vehicle (PG) or 4-NQO for one month using qPCR. n = 3. Two-tailed Student’s T test, *P ≤ 0.05. (**D**) Evaluation of *Cxcl9* expression levels in 4-NQO treated mice (1 month) injected with an IgG control or an IFNγ neutralizing antibody. N = 6. Two-tailed Student’s T test, ****P ≤ 0.0001. (**E**) Evaluation of *Cxcl9* expression levels in mice treated with vehicle (PG) or 4-NQO for one week using qPCR. n = 3. Two-tailed Student’s T test, *P ≤ 0.05. (**F**) CD45 immunofluorescence treated with vehicle (PG) or 4-NQO for one week, n = 3. Scale bar = 50 µM. (**G**) Schematic summarizing the generation of primary cultures of mouse esophageal epithelial cells using mice treated with vehicle (PG) or 4-NQO for one month. (**H**) Evaluation of *Cxcl9* expression levels using qPCR in mouse esophageal epithelial cells treated with PBS or IFNγ (50ng/mL) for 4h. n = 4. One-way analysis of variance followed by a Tukey multiple comparisons test, *P ≤ 0.05, ****P ≤ 0.0001. (**I**) Evaluation of *Cxcl9* expression levels using qPCR in mouse esophageal epithelial cells treated with PBS or IFNγ (50ng/mL), and with DMSO or JSH23 (10μg/mL) for 4h. n = 3. One-way analysis of variance followed by a Sidak multiple comparisons test, ****P ≤ 0.0001.

Since CXCL9 expression is known to be stimulated by the inflammatory marker IFNγ [36], we next analyzed *Ifnγ* expression levels in 4-NQO treated mice by qPCR. As shown in **Figure 6C**, we observed a significant increase in *Ifnγ* expression levels in one month 4-NQO treated mice compared to vehicle controls. To determine that increased *Ifnγ* was upstream of *Cxcl9* expression following 4-NQO treatment, we performed *in vivo* IFNγ neutralization in mice. We observed a significant decrease in *Cxcl9* expression in 4-NQO mice treated with a neutralizing antibody against IFNγ, compared to 4-NQO mice treated with an IgG control antibody (**Figure 6D**). To determine if the *Cxcl9* expression preceded T cell recruitment, we determined *Cxcl9* expression in mice treated with 4-NQO or vehicle control for one week and observed a significant increase in *Cxcl9* expression (**Figure 6E**), while immunofluorescence showed that barely any immune cells were recruited to the esophagus at this time point (**Figure 6F**).

To further investigate the connection between epithelial IKKβ signaling and the IFNγ/CXCL9 axis during early esophageal carcinogenesis, we generated primary cultures of esophageal epithelial cells from wildtype mice treated with 4-NQO or vehicle (**Figure 6G**). To determine if IFNγ regulates Cxcl9 expression, we treated these primary cultures with IFNγ or PBS for 4 hours. We observed that IFNγ treatment significantly increased Cxcl9 expression in both vehicle and 4-NQO primary cultures, with a notably higher expression in the 4-NQO-treated cultures (**Figure 6H**). To assess whether this Cxcl9 expression was dependent on the NFκB pathway, we treated 4-NQO primary cultures with and without IFNγ, and with and without the NFκB inhibitor JSH23. JSH23 significantly attenuated the IFNγ-induced increase in Cxcl9 expression, demonstrating that IFNγ induces Cxcl9 expression through the NFκB pathway (**Figure 6I**).

## Discussion

During carcinogenesis, the immune landscape evolves from an initial attempt to eliminate cancer cells to a progressively immunosuppressive environment in later stages [39]. This evolution is particularly challenging to study in ESCC due to late-stage detection, limiting access to early samples [1,3]. Single-cell transcriptomic analysis of ESCC immune cells reveals an inflammatory response with immune-suppressive populations contributing to immune escape and tumor progression [40]. Additionally, a single-cell atlas in a mouse model using the carcinogen 4-NQO highlights the role of immune suppression and chronic inflammation in promoting tumor development by creating a permissive microenvironment [29]. Therefore, further exploration of the inflammatory pathways driving the transition from a normal to a precancerous state in ESCC is needed.

Leveraging a well-established mouse model of stepwise carcinogenesis in ESCC [29,30], we explored the inflammatory pathways driving the transition from a normal to a precancerous state in the esophageal epithelium. We observed increased IKKβ/NFκB signaling, which regulates inflammatory responses by controlling pro-inflammatory genes and mediators [41]. Our findings demonstrate that in early esophageal squamous cell carcinogenesis, epithelial IKKβ signaling is crucial in regulating the immune cell landscape by modulating the expression of immune cell-recruiting molecules. These findings align with previous studies showing IKKβ/NFκB signaling in epithelial cells as a key regulator of the immune landscape through the control of inflammatory mediators and chemokines [42,43].

High levels of Th1 CD4+ T cells, along with a proinflammatory type 1 signature characterized by elevated *Ifnγ, Gzmb*, and *Tbx21*, have been reported in the early stages of esophageal carcinogenesis in 4-NQO-treated mice [29]. We found that the decreased recruitment of immune cells to the esophagus in 4-NQO/*Ikkβ*^*EEC-KO*^ mice was primarily due to reduced recruitment of CD3+, CD4+, CD8+, and regulatory T cells, accompanied by lower expression levels of the T cell-related gene *Cxcr3* and its chemoattractant *Cxcl9* [36]. Furthermore, we demonstrated that IFNγ, produced by CD8+ and Th1 T cells among others [36] regulates *Cxcl9* expression in esophageal epithelial cells. Thus, our data suggest that loss of epithelial IKKβ signaling attenuates the development of a proinflammatory type 1 response, preventing the amplification of T cell recruitment and the persistence of inflammation in the early stages of esophageal carcinogenesis.

Given that CD8+ T cells and Th1 CD4+ T cells are known to play crucial roles in killing tumor cells in many cancers [39], our observation of fewer recruited T cells and delayed cancer progression upon *Ikkβ* deletion might seem contradictory. However, a recent single-cell transcriptomic study showed a shift from a high type 1 inflammatory signature, to a type 3 inflammatory signature, with an increase in Th17 cells during the transition from hyperplasia to dysplasia in esophageal carcinogenesis [29]. Additionally, neutralizing IFNγ signaling in a mouse model of spontaneous chronic colorectal inflammation and cancer (*Socs1*-deficient mice) blocked the inflammatory response and cancer development [44]. Therefore, it is possible that the reduction of initial type 1 inflammation caused by the loss of epithelial IKKβ signaling prevented the transition to a protumorigenic type 3 inflammatory environment, thereby inhibiting esophageal cancer progression. Additionally, we observed changes in the overall immune landscape that may also contribute to decreased cancer progression. Future studies are needed to further characterize the changes in the immune landscape and inflammatory signatures following loss of epithelial IKKβ signaling in the early stages of esophageal carcinogenesis.

Despite the overall decrease in total immune cell recruitment, we observed a significant increase in macrophage recruitment to the esophageal epithelium, suggesting a role for these recruited macrophages in modulating the local inflammatory environment to delay esophageal carcinogenesis. Macrophages are highly plastic and can adopt diverse functional roles [45]. Traditionally, they have been classified as M1 (‘classically activated’ and proinflammatory) and M2 (‘alternatively activated,’ anti-inflammatory, and associated with tissue repair) [46]. However, macrophages can exhibit a wide range of phenotypes, suggesting that their classification should be more accurately viewed as a spectrum [45]. Our RNA-sequencing analyses revealed changes in gene expression related to pathways associated with wound healing and macrophage function. We observed increased expression of *Alox15* and *Mmp12* in 4-NQO/*Ikkβ*^*EEC-KO*^ mice compared to 4-NQO/controls. ALOX15 regulates inflammation through multiple mechanisms [32], acts as a tumor suppressor in colorectal cancer [47]. Reduced levels of *Alox15* have been reported in various cancers, including esophageal cancer [48]. MMP12 is predominantly expressed by macrophages, especially alternatively activated ones [33]. In inflammatory diseases, MMP12 acts as a potent anti-inflammatory agent by inhibiting complement in arthritis [49] and promoting the resolution of myocardial infarction [50]. It also cleaves IFNγ in autoimmune diseases, promoting resolution and suppressing proinflammatory macrophage activation [33]. This suggests that this increased recruitment of macrophages upon loss of epithelial IKKβ signaling in 4-NQO treated mice likely inhibits the chronic inflammation necessary for cancer progression and promotes the resolution of injuries caused by exposure to 4-NQO. Given the complexity of macrophage phenotypes, additional studies are crucial to further characterize these cells. This can be achieved using complementary techniques, such as isolating macrophages from esophageal tissue and employing advanced next-generation sequencing strategies like single-cell RNA sequencing. Additionally, macrophage depletion experiments will be necessary to determine the precise role of macrophages following the loss of epithelial *Ikkβ* in the context of esophageal precancer.

In summary, our research highlights the crucial role that epithelial IKKβ/NFκB signaling plays in the transition from normal to precancerous esophagus. This signaling pathway modulates the immune response by regulating T cell and macrophage recruitment, and IFNγ/CXCL9 signaling. Understanding the pathways regulating the early stages of esophageal carcinogenesis is essential for developing better methods to detect and treat esophageal squamous cell cancer at its early stages.

## Acknowledgements

We would like to thank Lia Tsikretsis for her assistance in animal care during the course of this study.

## Author Contributions

N.H., M-P.T. conceived and designed the experiments. N.H. performed the experiments and did the data analysis. N.H., M-P.T. drafted the manuscript. M-P.T. provided funding support. All authors participated in the revision of the manuscript and approved the submitted version.

## Ethics Statement

All animal studies were approved by the Institutional Animal Care and Use Committee at Northwestern University.

