## Supplemental Table 1 for "Epithelial *Ikkβ* deletion modulates immune responses and the IFN_*γ*_/CXCL9 axis during early esophageal carcinogenesis"

| **Supplementary Table 1. Genes differentially expressed in 4-NQO/*Ikkβ^EEC-KO^* mice compared to 4-NQO/control mice** | | | |
| --- | --- | --- | --- |
| **Gene Symbol** | **Log2 Fold Change** | **FDR Adjusted P Value** | **Description** |
| [2610528A11Rik](http://www.ncbi.nlm.nih.gov/gene/70045) | 1.483 | 2.24E-57 | RIKEN cDNA 2610528A11 gene |
| [Ces1b](http://www.ncbi.nlm.nih.gov/gene/382044) | -1.597 | 4.82E-50 | carboxylesterase 1B |
| [Slc6a2](http://www.ncbi.nlm.nih.gov/gene/20538) | -1.352 | 7.25E-32 | solute carrier family 6 (neurotransmitter transporter, noradrenalin), member 2 |
| [Tmprss11g](http://www.ncbi.nlm.nih.gov/gene/320454) | 1.580 | 8.76E-31 | transmembrane protease, serine 11g |
| [Fam171a2](http://www.ncbi.nlm.nih.gov/gene/217219) | 1.301 | 1.21E-28 | family with sequence similarity 171, member A2 |
| [Sprr2h](http://www.ncbi.nlm.nih.gov/gene/20762) | 1.866 | 1.21E-28 | small proline-rich protein 2H |
| [Nat8l](http://www.ncbi.nlm.nih.gov/gene/269642) | 1.713 | 1.21E-28 | N-acetyltransferase 8-like |
| [Ces1g](http://www.ncbi.nlm.nih.gov/gene/12623) | -1.196 | 1.93E-25 | carboxylesterase 1G |
| [Slc46a2](http://www.ncbi.nlm.nih.gov/gene/30936) | -1.420 | 1.03E-24 | solute carrier family 46, member 2 |
| [Skint3](http://www.ncbi.nlm.nih.gov/gene/195564) | 1.217 | 1.35E-23 | selection and upkeep of intraepithelial T cells 3 |
| [Cwh43](http://www.ncbi.nlm.nih.gov/gene/231293) | 1.292 | 3.41E-23 | cell wall biogenesis 43 C-terminal homolog |
| [Socs3](http://www.ncbi.nlm.nih.gov/gene/12702) | 1.169 | 1.34E-21 | suppressor of cytokine signaling 3 |
| [Urah](http://www.ncbi.nlm.nih.gov/gene/76974) | 1.544 | 3.02E-21 | urate (5-hydroxyiso-) hydrolase |
| [Elf3](http://www.ncbi.nlm.nih.gov/gene/13710) | 1.369 | 6.66E-19 | E74-like factor 3 |
| [Gm1110](http://www.ncbi.nlm.nih.gov/gene/382064) | -1.513 | 2.83E-18 | predicted gene 1110 |
| [Cyp2a5](http://www.ncbi.nlm.nih.gov/gene/13087) | -4.309 | 6.36E-17 | cytochrome P450, family 2, subfamily a, polypeptide 5 |
| [Dsg3](http://www.ncbi.nlm.nih.gov/gene/13512) | 1.172 | 6.50E-17 | desmoglein 3 |
| [Slc1a3](http://www.ncbi.nlm.nih.gov/gene/20512) | -1.637 | 4.22E-16 | solute carrier family 1 (glial high affinity glutamate transporter), member 3 |
| [Sprr2j-ps](http://www.ncbi.nlm.nih.gov/gene/20764) | 1.119 | 9.16E-16 | small proline-rich protein 2J, pseudogene |
| [Anxa8](http://www.ncbi.nlm.nih.gov/gene/11752) | 1.030 | 3.06E-15 | annexin A8 |
| [Gstm1](http://www.ncbi.nlm.nih.gov/gene/14862) | -1.085 | 3.06E-15 | glutathione S-transferase, mu 1 |
| [Ly6c1](http://www.ncbi.nlm.nih.gov/gene/17067) | 1.187 | 3.13E-15 | lymphocyte antigen 6 complex, locus C1 |
| [Mt3](http://www.ncbi.nlm.nih.gov/gene/17751) | -1.310 | 6.78E-15 | metallothionein 3 |
| [Tsku](http://www.ncbi.nlm.nih.gov/gene/244152) | -1.001 | 1.29E-14 | tsukushi, small leucine rich proteoglycan |
| [Prr9](http://www.ncbi.nlm.nih.gov/gene/109314) | -1.169 | 1.40E-14 | proline rich 9 |
| [Aldh3a1](http://www.ncbi.nlm.nih.gov/gene/11670) | -1.189 | 8.28E-14 | aldehyde dehydrogenase family 3, subfamily A1 |
| [Bcl3](http://www.ncbi.nlm.nih.gov/gene/12051) | 1.111 | 1.64E-13 | B cell leukemia/lymphoma 3 |
| [Selenbp1](http://www.ncbi.nlm.nih.gov/gene/20341) | -1.031 | 1.86E-13 | selenium binding protein 1 |
| [Mmp13](http://www.ncbi.nlm.nih.gov/gene/17386) | 2.237 | 2.96E-13 | matrix metallopeptidase 13 |
| [Cpn1](http://www.ncbi.nlm.nih.gov/gene/93721) | 1.106 | 6.09E-13 | carboxypeptidase N, polypeptide 1 |
| [Sprr2a3](http://www.ncbi.nlm.nih.gov/gene/100042514) | 3.525 | 1.12E-11 | small proline-rich protein 2A3 |
| [Klk6](http://www.ncbi.nlm.nih.gov/gene/19144) | 1.289 | 1.62E-11 | kallikrein related-peptidase 6 |
| [Grip2](http://www.ncbi.nlm.nih.gov/gene/243547) | 2.196 | 1.88E-11 | glutamate receptor interacting protein 2 |
| [Arl4d](http://www.ncbi.nlm.nih.gov/gene/80981) | 1.404 | 1.88E-11 | ADP-ribosylation factor-like 4D |
| [Ifi44](http://www.ncbi.nlm.nih.gov/gene/99899) | -1.101 | 2.54E-11 | interferon-induced protein 44 |
| [Mmp12](http://www.ncbi.nlm.nih.gov/gene/17381) | 2.730 | 5.93E-11 | matrix metallopeptidase 12 |
| [Fmo9](http://www.ncbi.nlm.nih.gov/gene/240894) | -1.439 | 1.03E-10 | flavin containing monooxygenase 9 |
| [Gdap1l1](http://www.ncbi.nlm.nih.gov/gene/228858) | -1.652 | 1.25E-10 | ganglioside-induced differentiation-associated protein 1-like 1 |
| [2310002F09Rik](http://www.ncbi.nlm.nih.gov/gene/100504720) | -1.177 | 2.25E-10 | RIKEN cDNA 2310002F09 gene |
| [Trank1](http://www.ncbi.nlm.nih.gov/gene/320429) | -1.744 | 2.80E-10 | tetratricopeptide repeat and ankyrin repeat containing 1 |
| [Gng4](http://www.ncbi.nlm.nih.gov/gene/14706) | -2.206 | 3.52E-10 | guanine nucleotide binding protein (G protein), gamma 4 |
| [Fcgbp](http://www.ncbi.nlm.nih.gov/gene/215384) | -1.375 | 3.72E-10 | Fc fragment of IgG binding protein |
| [Anks6](http://www.ncbi.nlm.nih.gov/gene/75691) | -1.021 | 4.90E-10 | ankyrin repeat and sterile alpha motif domain containing 6 |
| [Pcbp3](http://www.ncbi.nlm.nih.gov/gene/59093) | 1.322 | 5.19E-10 | poly(rC) binding protein 3 |
| [Ifi27l2a](http://www.ncbi.nlm.nih.gov/gene/76933) | -1.489 | 1.70E-09 | interferon, alpha-inducible protein 27 like 2A |
| [S1pr3](http://www.ncbi.nlm.nih.gov/gene/13610) | 1.721 | 1.80E-09 | sphingosine-1-phosphate receptor 3 |
| [Tafa4](http://www.ncbi.nlm.nih.gov/gene/320701) | -2.580 | 1.97E-09 | TAFA chemokine like family member 4 |
| [Abo](http://www.ncbi.nlm.nih.gov/gene/80908) | -1.092 | 4.31E-09 | ABO blood group (transferase A, alpha 1-3-N-acetylgalactosaminyltransferase, transferase B, alpha 1-3-galactosyltransferase) |
| [Aldh3b3](http://www.ncbi.nlm.nih.gov/gene/73458) | 1.263 | 4.31E-09 | aldehyde dehydrogenase 3 family, member B3 |
| [Alox15](http://www.ncbi.nlm.nih.gov/gene/11687) | 4.756 | 4.31E-09 | arachidonate 15-lipoxygenase |
| [Sptbn5](http://www.ncbi.nlm.nih.gov/gene/640524) | -1.076 | 4.93E-09 | spectrin beta, non-erythrocytic 5 |
| [Adamts13](http://www.ncbi.nlm.nih.gov/gene/279028) | -1.697 | 8.09E-09 | a disintegrin-like and metallopeptidase (reprolysin type) with thrombospondin type 1 motif, 13 |
| [Stom](http://www.ncbi.nlm.nih.gov/gene/13830) | 1.536 | 9.19E-09 | stomatin |
| [9530059O14Rik](http://www.ncbi.nlm.nih.gov/gene/319626) | -2.735 | 1.21E-08 | RIKEN cDNA 9530059O14 gene |
| [Sh2d5](http://www.ncbi.nlm.nih.gov/gene/230863) | 1.120 | 1.92E-08 | SH2 domain containing 5 |
| [Sp9](http://www.ncbi.nlm.nih.gov/gene/381373) | -1.341 | 4.35E-08 | trans-acting transcription factor 9 |
| [Tubb2b](http://www.ncbi.nlm.nih.gov/gene/73710) | 1.226 | 4.64E-08 | tubulin, beta 2B class IIB |
| [Ankdd1a](http://www.ncbi.nlm.nih.gov/gene/330963) | -1.282 | 5.02E-08 | ankyrin repeat and death domain containing 1A |
| [Lelp1](http://www.ncbi.nlm.nih.gov/gene/69332) | -1.448 | 9.39E-08 | late cornified envelope-like proline-rich 1 |
| [Zan](http://www.ncbi.nlm.nih.gov/gene/22635) | 2.312 | 9.48E-08 | zonadhesin |
| [Otud7a](http://www.ncbi.nlm.nih.gov/gene/170711) | 1.919 | 9.60E-08 | OTU domain containing 7A |
| [Ccser1](http://www.ncbi.nlm.nih.gov/gene/232035) | -1.278 | 2.33E-07 | coiled-coil serine rich 1 |
| [Chil4](http://www.ncbi.nlm.nih.gov/gene/104183) | 2.658 | 3.71E-07 | chitinase-like 4 |
| [Pgbd5](http://www.ncbi.nlm.nih.gov/gene/209966) | -1.108 | 4.60E-07 | piggyBac transposable element derived 5 |
| [Cystm1](http://www.ncbi.nlm.nih.gov/gene/66060) | 1.372 | 4.69E-07 | cysteine-rich transmembrane module containing 1 |
| [Pou3f1](http://www.ncbi.nlm.nih.gov/gene/18991) | 1.001 | 5.53E-07 | POU domain, class 3, transcription factor 1 |
| [Ms4a10](http://www.ncbi.nlm.nih.gov/gene/69826) | 1.185 | 6.33E-07 | membrane-spanning 4-domains, subfamily A, member 10 |
| [Gsdmc](http://www.ncbi.nlm.nih.gov/gene/83492) | 1.245 | 6.90E-07 | gasdermin C |
| [Ache](http://www.ncbi.nlm.nih.gov/gene/11423) | -1.456 | 1.08E-06 | acetylcholinesterase |
| [Apol8](http://www.ncbi.nlm.nih.gov/gene/239552) | 1.103 | 1.24E-06 | apolipoprotein L 8 |
| [Usp13](http://www.ncbi.nlm.nih.gov/gene/72607) | -1.980 | 1.31E-06 | ubiquitin specific peptidase 13 (isopeptidase T-3) |
| [Stfa2](http://www.ncbi.nlm.nih.gov/gene/20862) | 1.784 | 1.85E-06 | stefin A2 |
| [Kng2](http://www.ncbi.nlm.nih.gov/gene/385643) | -1.264 | 2.57E-06 | kininogen 2 |
| [Gstm3](http://www.ncbi.nlm.nih.gov/gene/14864) | -1.616 | 2.62E-06 | glutathione S-transferase, mu 3 |
| [Rassf6](http://www.ncbi.nlm.nih.gov/gene/73246) | -1.136 | 3.61E-06 | Ras association (RalGDS/AF-6) domain family member 6 |
| [Scn5a](http://www.ncbi.nlm.nih.gov/gene/20271) | -1.721 | 4.18E-06 | sodium channel, voltage-gated, type V, alpha |
| [Gm5414](http://www.ncbi.nlm.nih.gov/gene/406223) | 2.246 | 4.33E-06 | predicted gene 5414 |
| [Clec7a](http://www.ncbi.nlm.nih.gov/gene/56644) | 1.117 | 6.41E-06 | C-type lectin domain family 7, member a |
| [Tmprss11b](http://www.ncbi.nlm.nih.gov/gene/319875) | 1.119 | 7.04E-06 | transmembrane protease, serine 11B |
| [Il19](http://www.ncbi.nlm.nih.gov/gene/329244) | 2.889 | 8.92E-06 | interleukin 19 |
| [Sprr2b](http://www.ncbi.nlm.nih.gov/gene/20756) | 2.051 | 9.51E-06 | small proline-rich protein 2B |
| [Tent5a](http://www.ncbi.nlm.nih.gov/gene/212943) | 1.406 | 9.65E-06 | terminal nucleotidyltransferase 5A |
| [Cstdc5](http://www.ncbi.nlm.nih.gov/gene/100034684) | 1.674 | 1.00E-05 | cystatin domain containing 5 |
| [Npl](http://www.ncbi.nlm.nih.gov/gene/74091) | 1.515 | 1.21E-05 | N-acetylneuraminate pyruvate lyase |
| [Prss12](http://www.ncbi.nlm.nih.gov/gene/19142) | 1.267 | 1.24E-05 | protease, serine 12 neurotrypsin (motopsin) |
| [Vipr1](http://www.ncbi.nlm.nih.gov/gene/22354) | -1.135 | 1.44E-05 | vasoactive intestinal peptide receptor 1 |
| [Card11](http://www.ncbi.nlm.nih.gov/gene/108723) | -1.145 | 1.68E-05 | caspase recruitment domain family, member 11 |
| [Smyd1](http://www.ncbi.nlm.nih.gov/gene/12180) | -2.388 | 1.95E-05 | SET and MYND domain containing 1 |
| [Sprr2i](http://www.ncbi.nlm.nih.gov/gene/20763) | 2.180 | 3.15E-05 | small proline-rich protein 2I |
| [Gck](http://www.ncbi.nlm.nih.gov/gene/103988) | -1.454 | 3.38E-05 | glucokinase |
| [Mcub](http://www.ncbi.nlm.nih.gov/gene/66815) | 1.138 | 3.52E-05 | mitochondrial calcium uniporter dominant negative beta subunit |
| [Steap4](http://www.ncbi.nlm.nih.gov/gene/117167) | 1.135 | 3.68E-05 | STEAP family member 4 |
| [Ankrd24](http://www.ncbi.nlm.nih.gov/gene/70615) | -1.207 | 4.97E-05 | ankyrin repeat domain 24 |
| [Lrrc3b](http://www.ncbi.nlm.nih.gov/gene/218763) | -2.747 | 6.47E-05 | leucine rich repeat containing 3B |
| [Dab1](http://www.ncbi.nlm.nih.gov/gene/13131) | -1.706 | 6.80E-05 | disabled 1 |
| [Cyp2f2](http://www.ncbi.nlm.nih.gov/gene/13107) | -1.855 | 6.89E-05 | cytochrome P450, family 2, subfamily f, polypeptide 2 |
| [Klhl33](http://www.ncbi.nlm.nih.gov/gene/546611) | -2.045 | 7.46E-05 | kelch-like 33 |
| [Sprr1a](http://www.ncbi.nlm.nih.gov/gene/20753) | 1.257 | 7.86E-05 | small proline-rich protein 1A |
| [Slurp2](http://www.ncbi.nlm.nih.gov/gene/69462) | -1.287 | 8.26E-05 | secreted Ly6/Plaur domain containing 2 |
| [Txlnb](http://www.ncbi.nlm.nih.gov/gene/378431) | -1.623 | 9.88E-05 | taxilin beta |
| [Sptssb](http://www.ncbi.nlm.nih.gov/gene/66183) | -1.065 | 9.88E-05 | serine palmitoyltransferase, small subunit B |
| [Tmeff2](http://www.ncbi.nlm.nih.gov/gene/56363) | -1.072 | 9.89E-05 | transmembrane protein with EGF-like and two follistatin-like domains 2 |
| [Ifi213](http://www.ncbi.nlm.nih.gov/gene/623121) | -1.082 | 0.000100046 | interferon activated gene 213 |
| [Cd36](http://www.ncbi.nlm.nih.gov/gene/12491) | -1.346 | 0.000102809 | CD36 molecule |
| [Catsperg1](http://www.ncbi.nlm.nih.gov/gene/320225) | -1.047 | 0.000104935 | cation channel sperm associated auxiliary subunit gamma 1 |
| [Krt6b](http://www.ncbi.nlm.nih.gov/gene/16688) | 1.730 | 0.000107242 | keratin 6B |
| [Gm5478](http://www.ncbi.nlm.nih.gov/gene/432987) | 1.524 | 0.000119235 | predicted pseudogene 5478 |
| [Sema3d](http://www.ncbi.nlm.nih.gov/gene/108151) | -1.142 | 0.000149791 | sema domain, immunoglobulin domain (Ig), short basic domain, secreted, (semaphorin) 3D |
| [Fos](http://www.ncbi.nlm.nih.gov/gene/14281) | 1.650 | 0.000150455 | FBJ osteosarcoma oncogene |
| [Bfsp1](http://www.ncbi.nlm.nih.gov/gene/12075) | 1.277 | 0.000159279 | beaded filament structural protein 1, in lens-CP94 |
| [Cadm4](http://www.ncbi.nlm.nih.gov/gene/260299) | 1.093 | 0.00016983 | cell adhesion molecule 4 |
| [Gm26797](http://www.ncbi.nlm.nih.gov/gene/108167690) | -1.153 | 0.00017048 | predicted gene, 26797 |
| [Lemd1](http://www.ncbi.nlm.nih.gov/gene/213409) | 1.101 | 0.000192659 | LEM domain containing 1 |
| [Aox3](http://www.ncbi.nlm.nih.gov/gene/71724) | -1.387 | 0.000194795 | aldehyde oxidase 3 |
| [Ces1d](http://www.ncbi.nlm.nih.gov/gene/104158) | -1.128 | 0.000195855 | carboxylesterase 1D |
| [Adm](http://www.ncbi.nlm.nih.gov/gene/11535) | -1.103 | 0.000212625 | adrenomedullin |
| [Apol9a](http://www.ncbi.nlm.nih.gov/gene/223672) | -1.369 | 0.000232485 | apolipoprotein L 9a |
| [Ntrk3](http://www.ncbi.nlm.nih.gov/gene/18213) | -1.084 | 0.00023589 | neurotrophic tyrosine kinase, receptor, type 3 |
| [E430016F16Rik](http://www.ncbi.nlm.nih.gov/gene/414121) | -1.084 | 0.00023589 | RIKEN cDNA E430016F16 gene |
| [Scgb1a1](http://www.ncbi.nlm.nih.gov/gene/22287) | -1.133 | 0.00025202 | secretoglobin, family 1A, member 1 (uteroglobin) |
| [Slc38a3](http://www.ncbi.nlm.nih.gov/gene/76257) | -1.246 | 0.000263861 | solute carrier family 38, member 3 |
| [F10](http://www.ncbi.nlm.nih.gov/gene/14058) | 1.960 | 0.000286919 | coagulation factor X |
| [Krt84](http://www.ncbi.nlm.nih.gov/gene/16680) | 1.305 | 0.000327179 | keratin 84 |
| [Kcnc4](http://www.ncbi.nlm.nih.gov/gene/99738) | 1.330 | 0.000366548 | potassium voltage gated channel, Shaw-related subfamily, member 4 |
| [P4ha3](http://www.ncbi.nlm.nih.gov/gene/320452) | 1.380 | 0.000388036 | procollagen-proline, 2-oxoglutarate 4-dioxygenase (proline 4-hydroxylase), alpha polypeptide III |
| [Ndufa4l2](http://www.ncbi.nlm.nih.gov/gene/407790) | 1.198 | 0.000426703 | Ndufa4, mitochondrial complex associated like 2 |
| [Tmem45b](http://www.ncbi.nlm.nih.gov/gene/235135) | 1.026 | 0.00042762 | transmembrane protein 45b |
| [Kmo](http://www.ncbi.nlm.nih.gov/gene/98256) | -1.366 | 0.000439662 | kynurenine 3-monooxygenase (kynurenine 3-hydroxylase) |
| [Nos1](http://www.ncbi.nlm.nih.gov/gene/18125) | -1.853 | 0.000461614 | nitric oxide synthase 1, neuronal |
| [Hbegf](http://www.ncbi.nlm.nih.gov/gene/15200) | 1.055 | 0.000483378 | heparin-binding EGF-like growth factor |
| [Spp1](http://www.ncbi.nlm.nih.gov/gene/20750) | 1.905 | 0.000554817 | secreted phosphoprotein 1 |
| [Adh6a](http://www.ncbi.nlm.nih.gov/gene/69117) | -1.539 | 0.000671558 | alcohol dehydrogenase 6A (class V) |
| [Lmod3](http://www.ncbi.nlm.nih.gov/gene/320502) | -2.819 | 0.000740221 | leiomodin 3 (fetal) |
| [Casq1](http://www.ncbi.nlm.nih.gov/gene/12372) | -1.942 | 0.000950359 | calsequestrin 1 |
| [Lmod2](http://www.ncbi.nlm.nih.gov/gene/93677) | -2.324 | 0.00097178 | leiomodin 2 (cardiac) |
| [Xkr6](http://www.ncbi.nlm.nih.gov/gene/219149) | -1.009 | 0.000992488 | X-linked Kx blood group related 6 |
| [Flt3](http://www.ncbi.nlm.nih.gov/gene/14255) | -1.199 | 0.000993829 | FMS-like tyrosine kinase 3 |
| [F7](http://www.ncbi.nlm.nih.gov/gene/14068) | 2.140 | 0.001037481 | coagulation factor VII |
| [Flg2](http://www.ncbi.nlm.nih.gov/gene/229574) | -1.144 | 0.001119325 | filaggrin family member 2 |
| [Rhpn1](http://www.ncbi.nlm.nih.gov/gene/14787) | -1.310 | 0.001194839 | rhophilin, Rho GTPase binding protein 1 |
| [Lingo2](http://www.ncbi.nlm.nih.gov/gene/242384) | -2.413 | 0.001248738 | leucine rich repeat and Ig domain containing 2 |
| [Chil3](http://www.ncbi.nlm.nih.gov/gene/12655) | 2.718 | 0.001360223 | chitinase-like 3 |
| [Dnah10](http://www.ncbi.nlm.nih.gov/gene/56087) | -1.181 | 0.0015192 | dynein, axonemal, heavy chain 10 |
| [Pkig](http://www.ncbi.nlm.nih.gov/gene/18769) | -1.014 | 0.001567192 | protein kinase inhibitor, gamma |
| [Morn4](http://www.ncbi.nlm.nih.gov/gene/226123) | -1.463 | 0.001581168 | MORN repeat containing 4 |
| [Coro6](http://www.ncbi.nlm.nih.gov/gene/216961) | -1.438 | 0.002039945 | coronin 6 |
| [Ces1e](http://www.ncbi.nlm.nih.gov/gene/13897) | -1.217 | 0.002127959 | carboxylesterase 1E |
| [Ahsg](http://www.ncbi.nlm.nih.gov/gene/11625) | -1.318 | 0.002332523 | alpha-2-HS-glycoprotein |
| [Myo18b](http://www.ncbi.nlm.nih.gov/gene/74376) | -1.133 | 0.002539667 | myosin XVIIIb |
| [Dynap](http://www.ncbi.nlm.nih.gov/gene/75577) | 1.328 | 0.002605785 | dynactin associated protein |
| [Defb3](http://www.ncbi.nlm.nih.gov/gene/27358) | 1.119 | 0.002660119 | defensin beta 3 |
| [Ifi206](http://www.ncbi.nlm.nih.gov/gene/102639543) | -1.039 | 0.002958621 | interferon activated gene 206 |
| [Ada](http://www.ncbi.nlm.nih.gov/gene/11486) | -1.662 | 0.002970489 | adenosine deaminase |
| [Tnfaip8l3](http://www.ncbi.nlm.nih.gov/gene/244882) | 1.251 | 0.003053851 | tumor necrosis factor, alpha-induced protein 8-like 3 |
| [Apol9b](http://www.ncbi.nlm.nih.gov/gene/71898) | -1.123 | 0.003102655 | apolipoprotein L 9b |
| [Prkcq](http://www.ncbi.nlm.nih.gov/gene/18761) | -1.265 | 0.003137368 | protein kinase C, theta |
| [Chga](http://www.ncbi.nlm.nih.gov/gene/12652) | -1.115 | 0.003444977 | chromogranin A |
| [5730420D15Rik](http://www.ncbi.nlm.nih.gov/gene/70523) | -1.868 | 0.003595705 | RIKEN cDNA 5730420D15 gene |
| [Acsm3](http://www.ncbi.nlm.nih.gov/gene/20216) | -1.065 | 0.003615683 | acyl-CoA synthetase medium-chain family member 3 |
| [A4gnt](http://www.ncbi.nlm.nih.gov/gene/333424) | -1.391 | 0.003756386 | alpha-1,4-N-acetylglucosaminyltransferase |
| [Ldb3](http://www.ncbi.nlm.nih.gov/gene/24131) | -1.585 | 0.003764475 | LIM domain binding 3 |
| [Krt6a](http://www.ncbi.nlm.nih.gov/gene/16687) | 1.423 | 0.003924799 | keratin 6A |
| [0610031O16Rik](http://www.ncbi.nlm.nih.gov/gene/68369) | -1.150 | 0.004009441 | RIKEN cDNA 0610031O16 gene |
| [Ctnnd2](http://www.ncbi.nlm.nih.gov/gene/18163) | -1.119 | 0.004112265 | catenin (cadherin associated protein), delta 2 |
| [Cyb5r2](http://www.ncbi.nlm.nih.gov/gene/320635) | -1.205 | 0.004369997 | cytochrome b5 reductase 2 |
| [Sprr1b](http://www.ncbi.nlm.nih.gov/gene/20754) | 1.154 | 0.00441508 | small proline-rich protein 1B |
| [Snhg11](http://www.ncbi.nlm.nih.gov/gene/319317) | -1.717 | 0.004530005 | small nucleolar RNA host gene 11 |
| [Gsdmc2](http://www.ncbi.nlm.nih.gov/gene/331063) | 1.250 | 0.004621736 | gasdermin C2 |
| [Clec9a](http://www.ncbi.nlm.nih.gov/gene/232414) | -1.437 | 0.004734325 | C-type lectin domain family 9, member a |
| [Lrtm1](http://www.ncbi.nlm.nih.gov/gene/319476) | -1.296 | 0.004944673 | leucine-rich repeats and transmembrane domains 1 |
| [Cxcr3](http://www.ncbi.nlm.nih.gov/gene/12766) | -1.279 | 0.005633867 | chemokine (C-X-C motif) receptor 3 |
| [Rp1l1](http://www.ncbi.nlm.nih.gov/gene/271209) | -1.289 | 0.005821485 | retinitis pigmentosa 1 homolog like 1 |
| [Dock3](http://www.ncbi.nlm.nih.gov/gene/208869) | -1.400 | 0.006120619 | dedicator of cyto-kinesis 3 |
| [Far2](http://www.ncbi.nlm.nih.gov/gene/330450) | -1.360 | 0.006188428 | fatty acyl CoA reductase 2 |
| [Krt16](http://www.ncbi.nlm.nih.gov/gene/16666) | 1.780 | 0.006194284 | keratin 16 |
| [Adam23](http://www.ncbi.nlm.nih.gov/gene/23792) | -1.003 | 0.00635378 | a disintegrin and metallopeptidase domain 23 |
| [Pilra](http://www.ncbi.nlm.nih.gov/gene/231805) | 1.067 | 0.006954515 | paired immunoglobin-like type 2 receptor alpha |
| [Kctd14](http://www.ncbi.nlm.nih.gov/gene/233529) | -1.294 | 0.007039579 | potassium channel tetramerisation domain containing 14 |
| [Arc](http://www.ncbi.nlm.nih.gov/gene/11838) | 1.200 | 0.007821901 | activity regulated cytoskeletal-associated protein |
| [Tespa1](http://www.ncbi.nlm.nih.gov/gene/67596) | -1.078 | 0.008029023 | thymocyte expressed, positive selection associated 1 |
| [Ptpn5](http://www.ncbi.nlm.nih.gov/gene/19259) | -1.174 | 0.008105832 | protein tyrosine phosphatase, non-receptor type 5 |
| [Tmprss6](http://www.ncbi.nlm.nih.gov/gene/71753) | -1.021 | 0.008292489 | transmembrane serine protease 6 |
| [Gm36595](http://www.ncbi.nlm.nih.gov/gene/102640560) | -1.186 | 0.008383906 | predicted gene, 36595 |
| [Nav3](http://www.ncbi.nlm.nih.gov/gene/260315) | -1.322 | 0.008420326 | neuron navigator 3 |
| [Sgk2](http://www.ncbi.nlm.nih.gov/gene/27219) | -1.824 | 0.008703089 | serum/glucocorticoid regulated kinase 2 |
| [Krt10](http://www.ncbi.nlm.nih.gov/gene/16661) | -1.576 | 0.008800128 | keratin 10 |
| [Lrrc73](http://www.ncbi.nlm.nih.gov/gene/224813) | -1.094 | 0.00957776 | leucine rich repeat containing 73 |
| [Pf4](http://www.ncbi.nlm.nih.gov/gene/56744) | 1.378 | 0.009912528 | platelet factor 4 |
| [Atp1b1](http://www.ncbi.nlm.nih.gov/gene/11931) | -1.016 | 0.010222179 | ATPase, Na+/K+ transporting, beta 1 polypeptide |
| [Fras1](http://www.ncbi.nlm.nih.gov/gene/231470) | -1.378 | 0.010274003 | Fraser extracellular matrix complex subunit 1 |
| [Sprr2d](http://www.ncbi.nlm.nih.gov/gene/20758) | 1.396 | 0.010336963 | small proline-rich protein 2D |
| [Nppb](http://www.ncbi.nlm.nih.gov/gene/18158) | 1.635 | 0.010438749 | natriuretic peptide type B |
| [Pi15](http://www.ncbi.nlm.nih.gov/gene/94227) | 1.105 | 0.010637472 | peptidase inhibitor 15 |
| [Pdzrn4](http://www.ncbi.nlm.nih.gov/gene/239618) | -1.390 | 0.010942418 | PDZ domain containing RING finger 4 |
| [Dsg4](http://www.ncbi.nlm.nih.gov/gene/16769) | 1.490 | 0.010984171 | desmoglein 4 |
| [Olah](http://www.ncbi.nlm.nih.gov/gene/99035) | -1.396 | 0.011600271 | oleoyl-ACP hydrolase |
| [Pygm](http://www.ncbi.nlm.nih.gov/gene/19309) | -1.723 | 0.01195731 | muscle glycogen phosphorylase |
| [Tmem117](http://www.ncbi.nlm.nih.gov/gene/320709) | -1.037 | 0.012230044 | transmembrane protein 117 |
| [Siglech](http://www.ncbi.nlm.nih.gov/gene/233274) | -1.258 | 0.012477372 | sialic acid binding Ig-like lectin H |
| [Abca4](http://www.ncbi.nlm.nih.gov/gene/11304) | -1.120 | 0.012647829 | ATP-binding cassette, sub-family A (ABC1), member 4 |
| [Sprr2g](http://www.ncbi.nlm.nih.gov/gene/20761) | 1.631 | 0.012824812 | small proline-rich protein 2G |
| [Oas1h](http://www.ncbi.nlm.nih.gov/gene/246729) | -1.119 | 0.012875436 | 2'-5' oligoadenylate synthetase 1H |
| [Btla](http://www.ncbi.nlm.nih.gov/gene/208154) | -1.315 | 0.013064813 | B and T lymphocyte associated |
| [Bhlha15](http://www.ncbi.nlm.nih.gov/gene/17341) | 1.174 | 0.013717122 | basic helix-loop-helix family, member a15 |
| [Tmem132b](http://www.ncbi.nlm.nih.gov/gene/208151) | -1.520 | 0.013735069 | transmembrane protein 132B |
| [Sectm1b](http://www.ncbi.nlm.nih.gov/gene/58210) | -1.415 | 0.014106825 | secreted and transmembrane 1B |
| [C330022C24Rik](http://www.ncbi.nlm.nih.gov/gene/78520) | 1.654 | 0.014862533 | RIKEN cDNA C330022C24 gene |
| [H2-M10.5](http://www.ncbi.nlm.nih.gov/gene/224761) | -1.490 | 0.014953134 | histocompatibility 2, M region locus 10.5 |
| [Pkhd1l1](http://www.ncbi.nlm.nih.gov/gene/192190) | -1.163 | 0.015194689 | polycystic kidney and hepatic disease 1-like 1 |
| [Aadacl4](http://www.ncbi.nlm.nih.gov/gene/435815) | -1.817 | 0.015199102 | arylacetamide deacetylase like 4 |
| [Wfdc21](http://www.ncbi.nlm.nih.gov/gene/66107) | -1.351 | 0.015698752 | WAP four-disulfide core domain 21 |
| [Cyp4f18](http://www.ncbi.nlm.nih.gov/gene/72054) | 1.086 | 0.015742802 | cytochrome P450, family 4, subfamily f, polypeptide 18 |
| [Kcng3](http://www.ncbi.nlm.nih.gov/gene/225030) | -1.526 | 0.015748807 | potassium voltage-gated channel, subfamily G, member 3 |
| [Nctc1](http://www.ncbi.nlm.nih.gov/gene/330677) | -1.799 | 0.017116756 | non-coding transcript 1 |
| [Mb](http://www.ncbi.nlm.nih.gov/gene/17189) | -2.038 | 0.017946308 | myoglobin |
| [Glis1](http://www.ncbi.nlm.nih.gov/gene/230587) | -1.152 | 0.018152849 | GLIS family zinc finger 1 |
| [Cxcl9](http://www.ncbi.nlm.nih.gov/gene/17329) | -1.148 | 0.018202654 | chemokine (C-X-C motif) ligand 9 |
| [D130043K22Rik](http://www.ncbi.nlm.nih.gov/gene/210108) | -1.278 | 0.018526055 | RIKEN cDNA D130043K22 gene |
| [Pld6](http://www.ncbi.nlm.nih.gov/gene/194908) | -1.171 | 0.018679831 | phospholipase D family, member 6 |
| [Gm15713](http://www.ncbi.nlm.nih.gov/gene/626693) | -1.282 | 0.018981815 | predicted gene 15713 |
| [Oas1e](http://www.ncbi.nlm.nih.gov/gene/231699) | 1.174 | 0.019897634 | 2'-5' oligoadenylate synthetase 1E |
| [H2-DMb2](http://www.ncbi.nlm.nih.gov/gene/15000) | -1.222 | 0.020113414 | histocompatibility 2, class II, locus Mb2 |
| [Treml2](http://www.ncbi.nlm.nih.gov/gene/328833) | -1.572 | 0.022220922 | triggering receptor expressed on myeloid cells-like 2 |
| [Csta2](http://www.ncbi.nlm.nih.gov/gene/76770) | 1.466 | 0.022538508 | cystatin A family member 2 |
| [Adamtsl2](http://www.ncbi.nlm.nih.gov/gene/77794) | -1.367 | 0.022839225 | ADAMTS-like 2 |
| [Nron](http://www.ncbi.nlm.nih.gov/gene/320482) | -1.167 | 0.023357781 | non-protein coding RNA, repressor of NFAT |
| [Cdkn2a](http://www.ncbi.nlm.nih.gov/gene/12578) | -1.478 | 0.023755663 | cyclin dependent kinase inhibitor 2A |
| [Cox7a1](http://www.ncbi.nlm.nih.gov/gene/12865) | -1.357 | 0.024779239 | cytochrome c oxidase subunit 7A1 |
| [Sidt1](http://www.ncbi.nlm.nih.gov/gene/320007) | -1.207 | 0.025049651 | SID1 transmembrane family, member 1 |
| [Sypl2](http://www.ncbi.nlm.nih.gov/gene/17306) | -1.883 | 0.02564428 | synaptophysin-like 2 |
| [Ces1f](http://www.ncbi.nlm.nih.gov/gene/234564) | -1.384 | 0.026026277 | carboxylesterase 1F |
| [Rragd](http://www.ncbi.nlm.nih.gov/gene/52187) | -1.059 | 0.026029864 | Ras-related GTP binding D |
| [Hemgn](http://www.ncbi.nlm.nih.gov/gene/93966) | -1.501 | 0.026996287 | hemogen |
| [Slc29a4](http://www.ncbi.nlm.nih.gov/gene/243328) | 1.387 | 0.027036416 | solute carrier family 29 (nucleoside transporters), member 4 |
| [Cd7](http://www.ncbi.nlm.nih.gov/gene/12516) | -1.239 | 0.028118474 | CD7 antigen |
| [Scart1](http://www.ncbi.nlm.nih.gov/gene/244233) | -1.460 | 0.028530018 | scavenger receptor family member expressed on T cells 1 |
| [Gpd1](http://www.ncbi.nlm.nih.gov/gene/14555) | -1.524 | 0.0290364 | glycerol-3-phosphate dehydrogenase 1 (soluble) |
| [Neb](http://www.ncbi.nlm.nih.gov/gene/17996) | -1.535 | 0.02925145 | nebulin |
| [Phospho1](http://www.ncbi.nlm.nih.gov/gene/237928) | -1.023 | 0.031381324 | phosphatase, orphan 1 |
| [4933400A11Rik](http://www.ncbi.nlm.nih.gov/gene/66747) | -1.151 | 0.031389939 | capping protein (actin filament) muscle Z-line, alpha 1 pseudogene |
| [Sarm1](http://www.ncbi.nlm.nih.gov/gene/237868) | -1.187 | 0.031771654 | sterile alpha and HEAT/Armadillo motif containing 1 |
| [3425401B19Rik](http://www.ncbi.nlm.nih.gov/gene/100504518) | -1.586 | 0.032893289 | RIKEN cDNA 3425401B19 gene |
| [1700097N02Rik](http://www.ncbi.nlm.nih.gov/gene/67522) | 1.145 | 0.034018849 | RIKEN cDNA 1700097N02 gene |
| [Stpg1](http://www.ncbi.nlm.nih.gov/gene/78806) | -1.409 | 0.034103262 | sperm tail PG rich repeat containing 1 |
| [Cmya5](http://www.ncbi.nlm.nih.gov/gene/76469) | -1.865 | 0.034763991 | cardiomyopathy associated 5 |
| [Alpk3](http://www.ncbi.nlm.nih.gov/gene/116904) | -1.512 | 0.03648242 | alpha-kinase 3 |
| [Ggt1](http://www.ncbi.nlm.nih.gov/gene/14598) | -1.161 | 0.037634449 | gamma-glutamyltransferase 1 |
| [Sell](http://www.ncbi.nlm.nih.gov/gene/20343) | -1.054 | 0.038569211 | selectin, lymphocyte |
| [Adm2](http://www.ncbi.nlm.nih.gov/gene/223780) | 1.309 | 0.038763024 | adrenomedullin 2 |
| [Skint1](http://www.ncbi.nlm.nih.gov/gene/639781) | 1.305 | 0.039296718 | selection and upkeep of intraepithelial T cells 1 |
| [Ajap1](http://www.ncbi.nlm.nih.gov/gene/230959) | -1.332 | 0.039576383 | adherens junction associated protein 1 |
| [1110002E22Rik](http://www.ncbi.nlm.nih.gov/gene/102634333) | -1.142 | 0.04096251 | RIKEN cDNA 1110002E22 gene |
| [Gm867](http://www.ncbi.nlm.nih.gov/gene/333670) | -1.361 | 0.041373245 | predicted gene 867 |
| [Krt75](http://www.ncbi.nlm.nih.gov/gene/109052) | 1.468 | 0.041674196 | keratin 75 |
| [Gm13032](http://www.ncbi.nlm.nih.gov/gene/100049161) | -1.090 | 0.041912777 | predicted gene 13032 |
| [Gm5577](http://www.ncbi.nlm.nih.gov/gene/434064) | -1.056 | 0.043604338 | predicted gene 5577 |
| [Gm8369](http://www.ncbi.nlm.nih.gov/gene/666926) | -1.323 | 0.046598981 | predicted gene 8369 |
| [Btbd16](http://www.ncbi.nlm.nih.gov/gene/330660) | -1.155 | 0.047859008 | BTB (POZ) domain containing 16 |
| [Tigd4](http://www.ncbi.nlm.nih.gov/gene/403175) | -1.081 | 0.048518445 | tigger transposable element derived 4 |
| [Krt1](http://www.ncbi.nlm.nih.gov/gene/16678) | -1.174 | 0.048757428 | keratin 1 |
| [Zfp385c](http://www.ncbi.nlm.nih.gov/gene/278304) | -1.207 | 0.049441289 | zinc finger protein 385C |
| [P2rx1](http://www.ncbi.nlm.nih.gov/gene/18436) | -1.378 | 0.049511032 | purinergic receptor P2X, ligand-gated ion channel, 1 |
